# Evolution of plasticity in production and transgenerational inheritance of small RNAs under dynamic environmental conditions

**DOI:** 10.1101/722363

**Authors:** Willian T.A.F. Silva, Sarah P. Otto, Simone Immler

## Abstract

In a changing environment, small RNAs (sRNAs) play an important role in the post-transcriptional regulation of gene expression and can vary in abundance depending on the conditions experienced by an individual (phenotypic plasticity) and its parents (non-genetic inheritance). Many sRNAs are unusual in that they can be produced in two ways, either using genomic DNA as the template (primary sRNAs) or existing sRNAs as the template (secondary sRNAs). Thus, organisms can evolve rapid plastic responses to their current environment by adjusting the amplification rate of sRNA templates. sRNA levels can also be transmitted transgenerationally by the direct transfer of either sRNAs or the proteins involved in amplification. Theory is needed to describe the selective forces acting on sRNA levels, accounting for the dual nature of sRNAs as regulatory elements and templates for amplification and for the potential to transmit sRNAs and their amplification agents to offspring. Here, we develop a model to study the dynamics of sRNA production and inheritance in a fluctuating environment. We tested the selective advantage of mutants capable of sRNA-mediated phenotypic plasticity within resident populations with fixed levels of sRNA transcription. Even when the resident was allowed to evolve an optimal constant rate of sRNA production, plastic amplification rates capable of responding to environmental conditions were favored. By contrast, mechanisms allowing sRNA transcripts or amplification agents to be inherited were favored primarily when parents and offspring face similar environments and when selection acts before the optimal level of sRNA can be reached within the organism. Our study provides a clear set of testable predictions for the evolution of sRNA-related mechanisms of phenotypic plasticity and transgenerational inheritance.

## Introduction

Faced with a continually changing environment, organisms have evolved to respond to their environment (phenotypic plasticity) and to transfer information about environmental conditions from parents to offspring (non-genetic inheritance), influencing a broad array of ecological and evolutionary processes [1; 2; 3; 4; 5; 6]. Several studies in animals and plants suggest that some non-genetic factors inherited across generations have the potential to be beneficial to both parents and their offspring [7; 8]. A key step in assessing the role of non-genetic inheritance in adaptive processes is to improve our understanding of the different mechanisms involved. Three important factors currently thought to play a role in phenotypic plasticity and non-genetic inheritance include DNA methylation, chromatin structure modifications, and certain families of RNAs [9; 2, we follow [1] in using “non-genetic” inheritance to encompass a broad array of mechanisms by which parents can influence offspring, including epigenetic inheritance]. In particular, small RNAs (sRNAs) have been shown to play a potentially important role in transferring information about parental conditions to offspring due to their diversity in function [10] and their apparently rapid evolution [11; 12]. Here, we develop a model to assess how changing environmental conditions drive the evolution of plasticity and inheritance of sRNA levels.

sRNAs are short (*<* 200 nucleotides) non-coding RNAs that have been recently discovered and described across the tree of life, from bacteria [13; 14; 11] to plants [15; 16; 17] and animals [18; 19; 20]. The functions vary for different sRNA families, but two key functions are relevant for their potential role in adaptive processes, namely the post-transcriptional regulation of gene expression [21; 22] and the preservation of genome integrity through the repression of parasitic DNA elements such as transposable elements (TEs) [23]. Two sRNA families that are particularly associated with these two functions are microRNAs (miRNA) and Piwi-interacting RNAs (piRNA) [24]. miRNAs are known to play a key role in post-transcriptional gene regulation and have been found to regulate adipocyte differentiation in humans [25], hematopoietic differentiation in house mice *Mus musculus* [26], cell proliferation and apoptosis in *Drosophila* [27; 28], the maternal-to-zygotic transition in zebrafish *Danio rerio* [29], and several other developmental processes [30; 31; 32; 33]. piRNAs on the other hand have been primarily associated with processes protecting the genome against TE activity, particularly in the germline [34; 35]. piRNAs have been implicated in the maintenance and protection of germ cells in the zebrafish [36], protection of germ cells against viral infections in domestic chicken *Gallus domesticus* [37], chromatin repression of genomic regions with actively transposing elements in the house mouse [23], differential piRNA expression in sperm as a response to dietary changes in rats *Rattus norvegicus* [38], as well as the maintenance of metabolic homeostasis and normal lifespan in *Drosophila* [39].

sRNA abundance is also known to respond to changes in environmental conditions. In *Drosophila*, for example, changes in ambient temperature result in drastic but reversible changes in composition and abundance of ovarian miRNAs and piRNAs, with inversely correlated changes in their predicted targets [40; 41]. Similarly, changes in temperature cause changes in miRNA expression in cotton *Gossypium sp.* [42] and rockcress *Arabidopsis* [43], and exposure to drought affects miRNA expression in tomato *Solanum lycopersicum* [44]. In the bacteria *Burkholderia thailandensis*, several environmental conditions such as pH, salt, antibiotics in addition to temperature have been shown to cause differential sRNA expression [45; 46].

Recent evidence suggests that these changes in sRNA expression levels may not only affect one generation but may be transferred across several subsequent generations [47; 7]. In the nematode *Caenorhabditis elegans*, for example, viral infections led to a parental response that transformed viral double-stranded RNA into siRNAs resulting in viral immunity. These siRNAs were transmitted from parents to offspring even in the absence of the initial stimulus and provided protection against the virus for up to three subsequent generations [48]. Further studies in *C. elegans* showed that starvation in one generation resulted in the transmission of sRNA-induced silencing of nutrition-related genes across three generations without further starvation [49], and nematodes exposed to increased temperature exhibited a differential sRNA-mediated gene-silencing response [50].

The magnitude of the sRNA response to environmental conditions and its transgenerational inheritance can depend on the duration of the stimulus [temperature stress, 47], indicating that sRNA levels may be responsive to the magnitude of environmental stress. The persistence of sRNAs across generations differs among sRNA families, but two characteristics facilitate sRNA inheritance: the ability to avoid removal by sRNA-degradation agents [51; 52] and the ability to template their own synthesis via amplification cycles [53]. The sensitivity of sRNA production to environmental factors and the potential for transmission of sRNAs over several generations suggest that sRNAs can play an important role in adaptation.

The production of sRNA depends on two main processes, sRNA transcription from genomic DNA (generating primary sRNAs) and amplification from sRNA templates (generating secondary sRNAs), which can either directly act on their target transcripts or participate in a feed-forward mechanism of amplification to produce new secondary sRNAs [34]. This genome-independent amplification mechanism is particularly important for piRNAs and small-interfering RNAs (siRNAs) [54; 34]. Several proteins, including members of the Argonaute protein family [e.g., Argo and Piwi, 55; 56] and RNA-dependent RNA polymerases [RdRPs, 57; 58] act as amplification agents by using secondary sRNA molecules as templates for the production of more sRNAs in amplification cycles known as the ping-pong cycle for piRNA and the RdRP-driven amplification cycle for siRNA.

sRNA levels, as well as the rate of amplification, can be transmitted from cell to cell. Within an individual generation, existing sRNAs are passed on to daughter cells, where they may be subsequently amplified [59; 54; 60]. Across generations, both sRNA molecules and the amplification agents (either the proteins themselves or the regulatory state of the corresponding genes) can be transmitted from parents to offspring [61; 48; 62; 63], providing a mechanism that allows non-genetic inheritance of parental conditions [64; 65].

Here, we develop a model to investigate how sRNA regulation and non-genetic inheritance evolve in a varying environment, thereby mediating plastic and transgenerational responses to the environment. Given the complexity of sRNA responses, the model allows both the primary production and secondary amplification of sRNAs, incorporates the effects of sRNA production on fitness as a function of the environment, considers the transfer of sRNA transcripts and amplification machinery across generations, and incorporates potential costs of sRNAs. Our results indicate when different strategies of sRNA production and inheritance (as shown in Figure 1) are expected to invade a resident wild type population where sRNA production and/or transfer rates are genetically fixed, providing a framework to understand the evolution of sRNAs as important mediators of environmental conditions.

**Figure 1:**
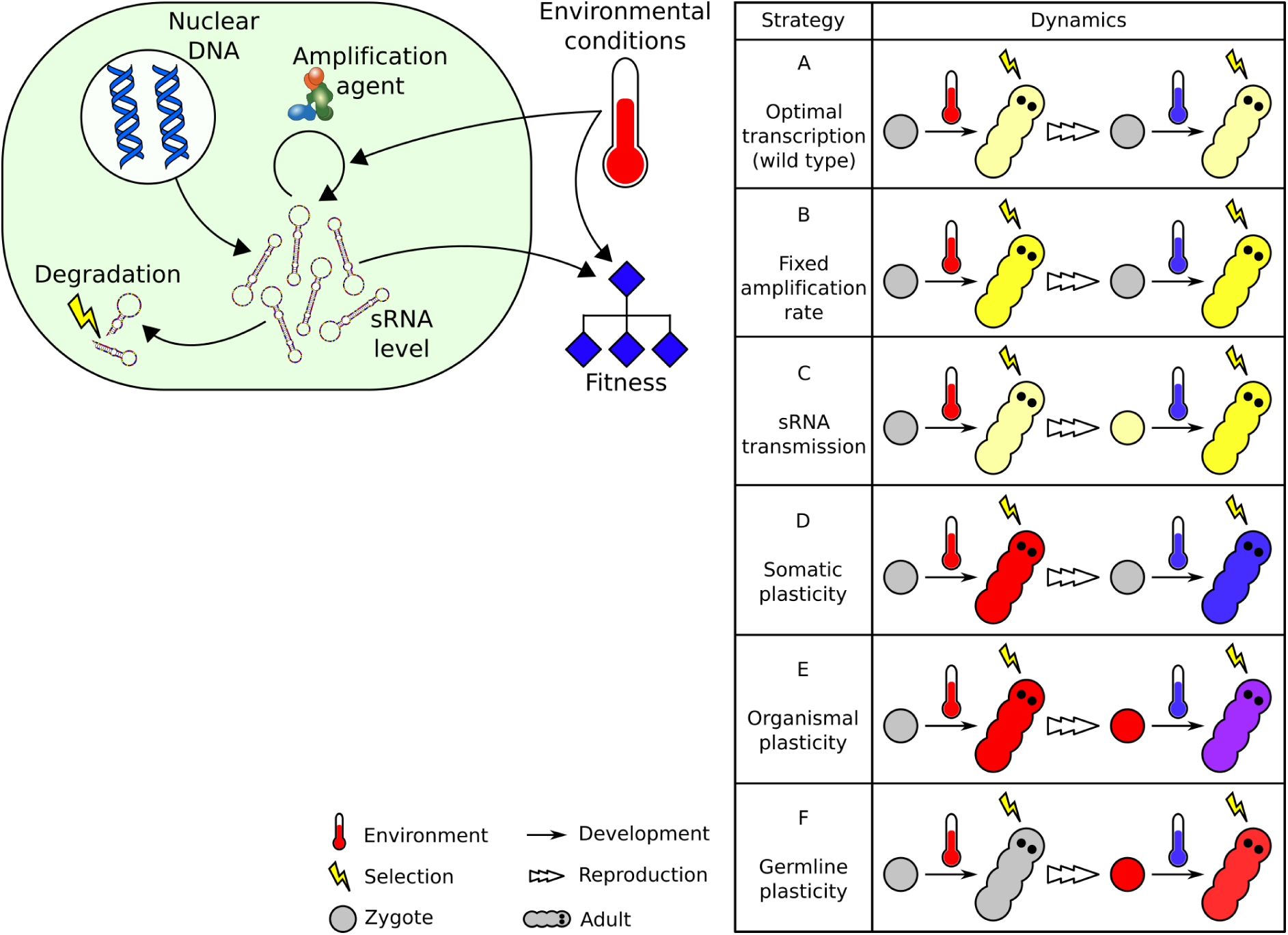
Model design. (A) sRNA production is a function of nuclear transcription, amplification and degradation processes. Fitness is a function of sRNA density and environmental conditions. (B) Strategies in the production and transmission of sRNA level or amplification rate that were explored in the simulations. Complete description in the main text.

## Methods

We developed a model that consists of three connected functions: (i) a function for the production, decay, and transmission of sRNA transcripts at the cellular level, (ii) a function to describe the amplification rate of sRNA and its inheritance, and (iii) a fitness function that depends on the costs and benefits of sRNA production in a given environment (Figure 1, left).

### sRNA production

sRNA production in our model is based on a feed-forward mechanism whereby the cellular level of sRNA (*n*, Equation 1) changes as a function of the transcription rate from nuclear DNA *µ*, the birth rate of sRNAs from existing template sRNAs, and the degradation rate *d*. Specifically, the total birth rate depends on the amplification rate *b* and on the number of existing templates (*n*), rising linearly with *n* when templates are rare but approaching a maximum amplification rate of *m* when *n* is high:

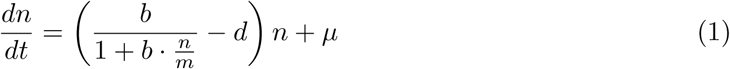

Equation 1 has two equilibria, only one of which is positive and is the biologically relevant equilibrium:

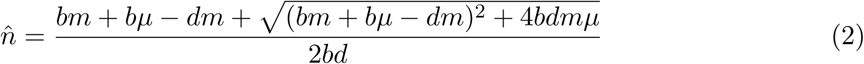

Time in Equation 1 is measured in cell generations. We assume that individuals replicate every *c* cell generations and set *c* = 20, as a default value. This lies within the range of cell divisions per generation within the germline reported for different animal species [66]: 8.5 in *C. elegans*, 36 in *D. melanogaster*, and 200 in humans.

Additionally, sRNA transcripts can be directly transmitted across generations through the gametes of the parents, and the sRNA amplification rate can change in response to current or past environmental conditions (*ϵ*). We therefore distinguished between the amplification rate in the soma, *b*\*, and the amplification rate in the germline, *b*^#^. We then defined six possible strategies for the production, responsiveness and transmission of sRNAs and their amplification rate (Figure 1, right):

A. Optimal transcription (wild type): Individuals have a fixed genetically determined sRNA production via transcription only, regardless of environmental conditions. There is no amplification (*b*\* = *b*^#^ = 0), plasticity or transgenerational transmission of sRNAs. Transcription rates are assumed to have evolved to their optimal levels.
B. Fixed amplification rate: In addition to the optimal transcription rate in strategy A, strategy B individuals have a fixed genetically determined amplification rate (*b*\* = *b*^#^ > 0), without the ability to transmit sRNA levels or respond to environmental conditions.
C. sRNA transmission: Similarly to strategy A, production of sRNA occurs via a fixed optimal transcription rate, but strategy C individuals can transmit sRNA levels to their offspring through their gametes.
D. Somatic plasticity: Individuals have a plastic somatic amplification rate (*b*\*) and a genetically fixed germline amplification rate (*b*^#^ = 0), allowing for a plastic response to changing environments in the parents without consequences for the next generation. In other words, in every generation, the somatic amplification rate *b*\* is reset to the inherited basal germline amplification rate *b*^#^.
E. Organismal plasticity: Individuals have a plastic somatic amplification rate *b*\* and a plastic germline amplification rate *b*^#^, which change in response to current environmental conditions, and these changes are transmitted to the next generation. Changes are global, so both the soma and the germline are affected equally (*b*\* = *b*^#^).
F. Germline plasticity: Individuals have a genetically fixed somatic amplification rate *b*\* and plastic germline amplification rate *b*^#^, which changes in response to current environmental conditions and is transmitted to the next generation and affects the somatic phenotype of the offspring. In this strategy, individuals do not directly benefit from having a plastic amplification rate but can potentially benefit their offspring by providing information about the parental environmental conditions.

Given a set of fixed biological parameter values (transcription rate, degradation rate and amplification activity), there is a value of the amplification rate *b*, called *b*_*Wmax,g*_, that results in the highest fitness for a particular environmental condition *ϵ*. We assume that an individual can only adjust its amplification rate based on the information about current environmental conditions (no information about long-term environmental dynamics) and that plasticity has evolved to be adaptive, bringing the value of *b* closer to the optimal *b*_*Wmax,g*_ in that generation. Plasticity is, however, limited and only allows the amplification rate to shift a fraction *P*_*b*_ of the way between the individual’s initial amplification rate *b* (inherited from the previous generation) and the optimum *b*_*Wmax,g*_. Plasticity is absent in strategies A-C (*P*_*b*_ = 0), which means that *b*\* = *b*^#^ is constant across individual generations:

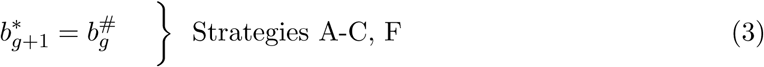

Somatic plasticity within an individual generation occurs for strategies D and E, such that the somatic amplification rate 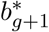 in generation *g* +1 differs from the adult germline in the previous generation, 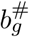:

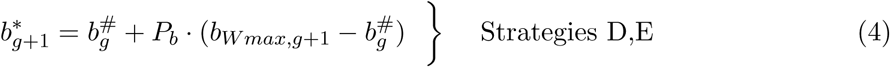

Transgenerational inheritance of the amplification rate, via transmission of templates, amplification agents, or epigenetic modification of those agents [67; 68; 69], is allowed in strategies E and F and is modeled via changes in the germline value of *b*^#^ from the adult parent (generation *g*) to adult offspring (generation *g* + 1), depending on the environment experienced between these two stages:

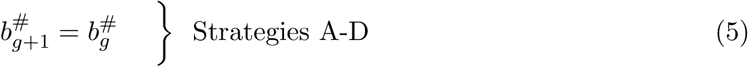

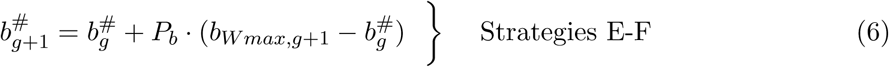

### Transmission rate

sRNAs in the germline can be transmitted through the gametes from parents to their off-spring [48; 62; 63]. This transmission, which distinguishes strategy C from the wildtype strategy A, provides a second opportunity for transgenerational inheritance of sRNAs, beyond the in-heritance of plastic changes in amplification rate. To model the transmission rate from parents to offspring, we vary the fraction *r*_*germ*_ of sRNAs that could be passed from the parental germ cell to the zygote. Specifically, given the density of sRNAs in adult cells (*n*_*final*_), the amount inherited in the zygote (*n*_*initial*_) in the next generation is set to *n*_*initial,g*+1_ = *r*_*germ*_ *· n*_*final,g*_. We assume that *n*_*final*_ results from the production of sRNAs throughout development according to Equation 1 and is the same in the germline and in somatic cells. In addition to being a characteristic of Strategy C, we briefly explore the evolution of the transmission ratio *r*_*germ*_ for strategies D-F.

### Environmental conditions

sRNA production is known to respond to a variety of environmental variables [45; 49; 43; 46; 42; 40; 41]. To model a response to the environment, we include a continuous variable *E* measuring the impact of the environment on individual fitness {*ϵ* ∈ ℝ : 0 ≤ *ϵ* ≤ 1}, where detrimental conditions (*ϵ* > 0) are deviations from the optimal environmental condition (*ϵ* = 0). For ease of reference, *ϵ* will represent a measure of how stressful the environment is to the organism. In benign environments (*ϵ* = 0), maximum fitness is achieved even when producing low to no amounts of the particular sRNA being modeled. In stressful environments, maximum fitness is achieved only by producing high amounts of this sRNA.

Transgenerational inheritance is predicted to be favorable when environmental conditions are more similar between parents and offspring than between more distant points in time (i.e., positively autocorrelated). In order to explore the impact of environmental autocorrelations, we hold the fraction of time that the environment was stressful (*ϵ* = 0.9) or benign (*ϵ* = 0.1) constant at 50% each 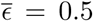. Thus, organisms always face the same array of environments, just in different orders over time. For different orderings, we use the probability that the environment is the same for parents and offspring, *p*_*ϵ*_ = 1 − [*k/*(*G* − 1)], as a measure of the environmental similarity of parents and offspring, where *k* is the total number of changes in environmental conditions (changes from *ϵ* = 0.1 to *ϵ* = 0.9 or vice-versa) and *G* is the total number of individual generations in each environmental scenario cycle (*G* = 20). High environmental similarity (high *p*_*ϵ*_) indicates that the environment switches rarely (positively autocorrelated), while a value of zero indicates switching every generation (negatively autocorrelated). Specifically, we consider environmental scenarios with *p*_*ϵ*_ ∈ {0.11, 0.53, 0.89} (see examples in Figure S1). For tractability, we assume that a given environmental scenario repeats every *G* = 20 individual generations, so that we could determine the long-term fitness of each strategy.

### Fitness

Individual fitness is modeled as a continuous function *W* (*n, E, P*_*b*_) of the amount of sRNA in the adult (*n*) after *c* = 20 cell generations, the current environmental condition (*ϵ*) and the degree of plasticity in amplification rate (*P*_*b*_; Equation 7):

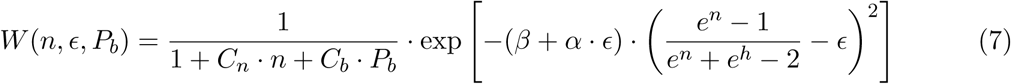

where the first term represents the cost (*C*_*n*_) associated with sRNA production (*n*) plus the cost associated with the plastic response in amplification rate *b* (*C*_*b*_). The second term of the fitness function is the benefit of matching sRNA production to environmental conditions. Specifically, environmental stress causes an exponentially declining fitness (with exponent *β* + *α ·ϵ*), which is ameliorated by a match between phenotype and the environment, *E*. The phenotype is a logistic-shaped function of the amount of sRNA, *n*, equaling 0 in the absence of sRNA and rising with large amounts of sRNA to 1, representing the best match to the most stressful environment (*ϵ* = 1). The parameter *h* represents the amount of sRNA required to match an intermediate environmental stress of *ϵ* = 0.5.

The form of equation 7 allows asymmetries in the fitness effects of sRNA production in stressful and benign environments (Figure 2). Specifically, the production of sRNAs in benign conditions is assumed to have a milder fitness effect than the benefits of production under stressful conditions. A plastic amplification rate (*P*_*b*_ > 0) has the benefit of bringing sRNA amounts closer to the optimum under different environmental conditions and can strongly increase fitness, but this plasticity in amplification rate comes at a cost (*C*_*b*_ *· P*_*b*_).

**Figure 2:**
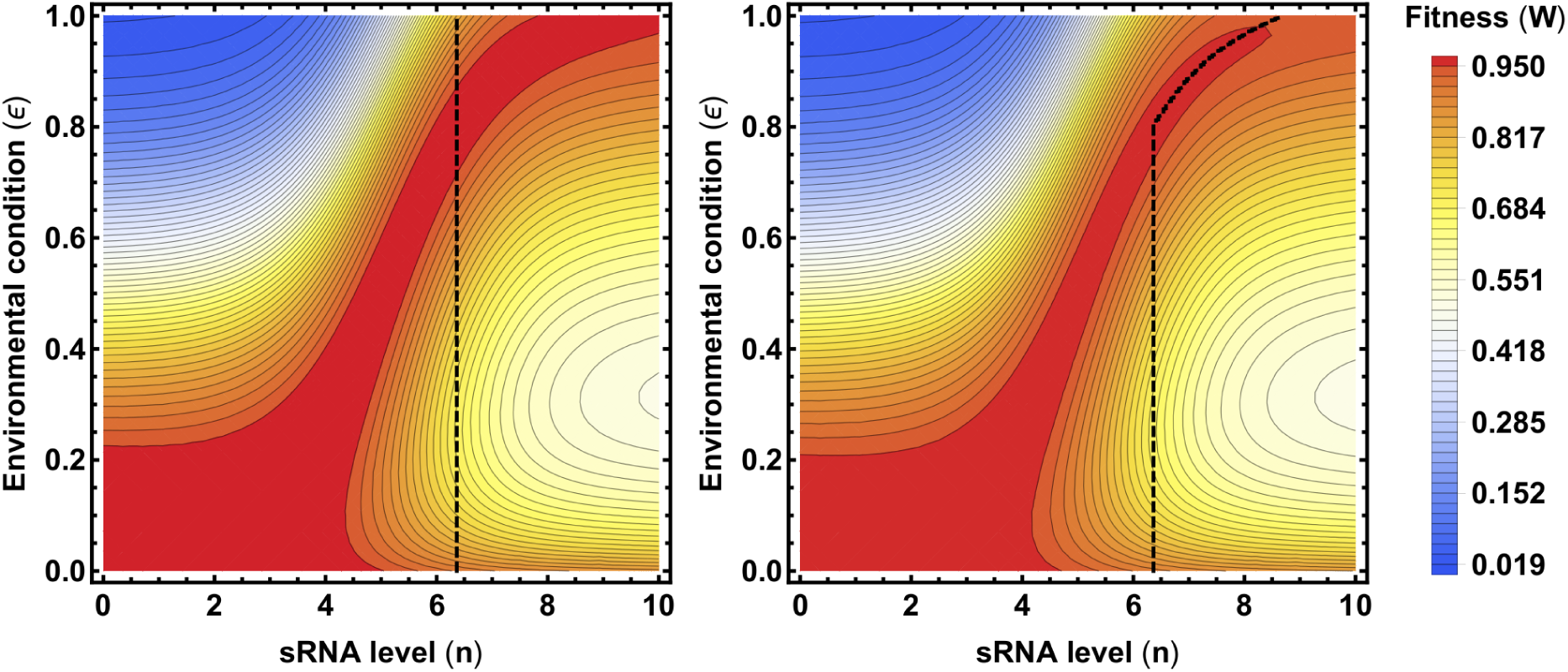
Fitness landscape as a function of the sRNA amount produced by the adult individual (*n*) and the environmental conditions (*ϵ*) with no plasticity in amplification rate (*P*_*b*_ = 0; left panel) and with maximum plasticity (*P*_*b*_ = 1.0; right panel). The dashed line on the left shows the constant amount of sRNAs produced by the wild type (strategy A, *P*_*b*_ = 0), and the dashed line on the right shows the amount of sRNAs produced by a plastic mutant (*P*_*b*_ = 1.0) that is able to amplify sRNA in response to stress. The following values in the fitness function were used across the study: *α* = 4.0, *β* = 0.1, *h* = 5.0, *C*_*n*_ = 0.005 and *C*_*b*_ = 2*C*_*n*_.

We assume a simple genetic model where each strategy can be encoded by a mutation at a single gene, in which case a mutant strategy will spread within a wildtype population if it has a higher geometric mean fitness (*W*_*GEO*_) over the set of environments encountered, where:

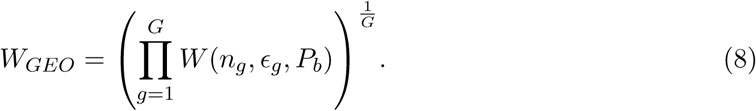

With sRNA transmission across generations (*r*_*germ*_ > 0), the amount of sRNAs in the zygote, *n*_*initial*_, depends on ancestral production of sRNA and varies across the repeated *G* = 20 periods; we thus calculated the geometric mean fitness after allowing *n*_*initial*_ at the beginning of a period to reach a stable value.

We next determine the environmental conditions under which mutant strategies (strategies B-F; superscript “+”) can invade a resident wildtype population (strategy A; superscript “−”), with parameters summarized in Table 1. To assess the strength of selection, on average, in favor of a mutant strategy, we use the selection coefficient *s*^+^ calculated from the relative geometric mean fitness of the mutant:

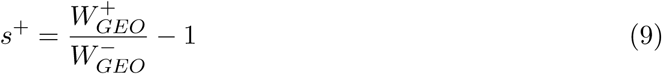

**Table 1:** Parameter values used for each strategy in the invasion analyses. S and G indicate plasticity in the soma and germline, respectively. Values in bold indicate mutant values.

| **Strategy** | $b$ | $d$ | $m$ | $\mu$ | $r_{germ}$ | $P_b$ |
| --- | --- | --- | --- | --- | --- | --- |
| *A* | 0 | 0.1 | 5.0 | 0.736 | 0 | 0 |
| **B** | **>0** | 0.1 | 5.0 | 0.736 | 0 | 0 |
| **C** | 0 | 0.1 | 5.0 | 0.736 | **>0** | 0 |
| **D** | **>0** | 0.1 | 5.0 | 0.736 | 0, **>0** | **S** |
| **E** | **>0** | 0.1 | 5.0 | 0.736 | 0, **>0** | **S+G** |
| **F** | **>0** | 0.1 | 5.0 | 0.736 | 0, **>0** | **G** |

Because we are particularly interested in cases where selection is strong, we illustrate where selection on the mutant becomes weaker in magnitude than | *s*^+^ |= 0.001, as drift is then likely to dominate selection even in populations of moderate size (*N* ≤ 1000; [70])

We assume that the degradation rate *d* is an intrinsic stability property of the sRNA molecule under focus and set it to *d*^−^ = *d*^+^ = 0.1. Similarly, the maximum amplification rate, *m*, is set to *m*^−^ = *m*^+^ = 5. We further assume that the rate of sRNA transcription from DNA has evolved to maximize the geometric mean fitness of strategy A in the absence of plasticity or transgenerational inheritance, setting this optimum parameter value as the default (*µ*^−^ = *µ*^+^ = 0.736 for the environmental conditions used in our simulations with 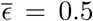 and *ϵ* ∈ {0.1, 0.9}). For strategies allowing sRNA amplification (B, D-F), we consider all positive values of *b*, assuming the value yielding the highest geometric mean fitness would eventually arise (*b*_*W max,g*_) and using this value to assess whether the mutant strategy could invade.

All analyses and simulations were performed in Wolfram Mathematica 11, version 11.0.1.0. A Mathematica notebook containing all of the analyses is available on the Dryad Digital Repository.

## Results

During individual development, the production of sRNA rises, approaching the steady state value, 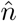 (Equation 2), at a rate that depends on the sRNA strategy (Table 1) and the parameters responsible for sRNA production and maintenance (*b, d, m*, and *µ*; Figure S2). The level of sRNA reached in adulthood, where we assess fitness, thus depends on the number of cell generations per individual, *c*, as well as the initial fraction of sRNA inherited by the zygote, *r*_*germ*_. The transmission ratio (*r*_*germ*_) determines how much sRNA levels are reset from generation to generation and hence the degree of sRNA oscillations witnessed across individual generations (Figure 3).

**Figure 3:**
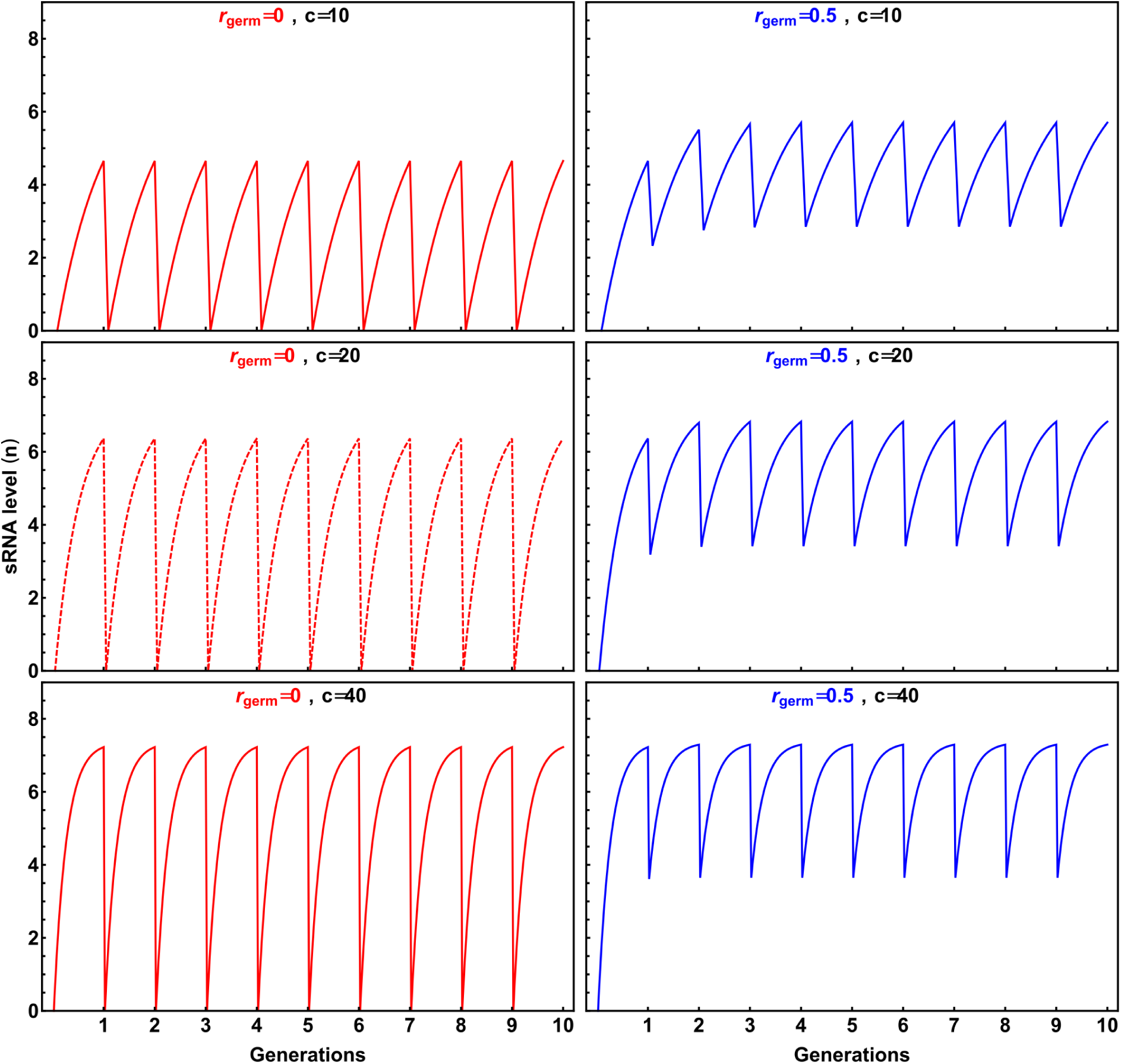
Sensitivity of the model with regard to the transgenerational transmission ratio (*r*_*germ*_; left) and number of cell divisions per generation (*c*; right). Red dashed curves represent the dynamics of sRNA production in strategy A. Parameter values used in the example: *b* = 0, *d* = 0.1, *m* = 5.0, *µ* = 0.736.

Without plasticity or non-genetic inheritance (*b* = 0, *r*_*germ*_ = 0, and *P*_*b*_ = 0), the fitness of strategy A is maximized at a transcription rate of *µ* = 0.736 (Figure S3). The adult level of sRNA then reached 6.36 for strategy A (solving Equation 1 at *c* = 20), a compromise between the optimal level in a benign environment (0) and in a stressful environment (8.68; Figure 2). Because strategy A lacks plasticity, the optimal transcription rate *µ* is the same regardless of the order of environments encountered. The dynamics of sRNA within and across individual generations for strategy A is shown in Figure 3 (dashed red lines).

### Evolution of mechanisms for sRNA amplification and inheritance

Mutants capable of amplifying sRNA production via a fixed amplification rate (*b*^+^ > 0, strategy are unable to invade a resident population of strategy A (i.e., *s*^+^ *<* 0) under the environmental conditions to which the resident population is already optimally adapted (*µ* = 0.736). Similarly, mutants capable of transmitting sRNA across generations (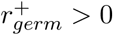, strategy C) are unable to invade a resident population of strategy A (Figure 4, upper panels). As neither of these strategies involve plastic responses to the environment, the results are insensitive to the order of environments encountered (*p*_*ϵ*_).

**Figure 4:**
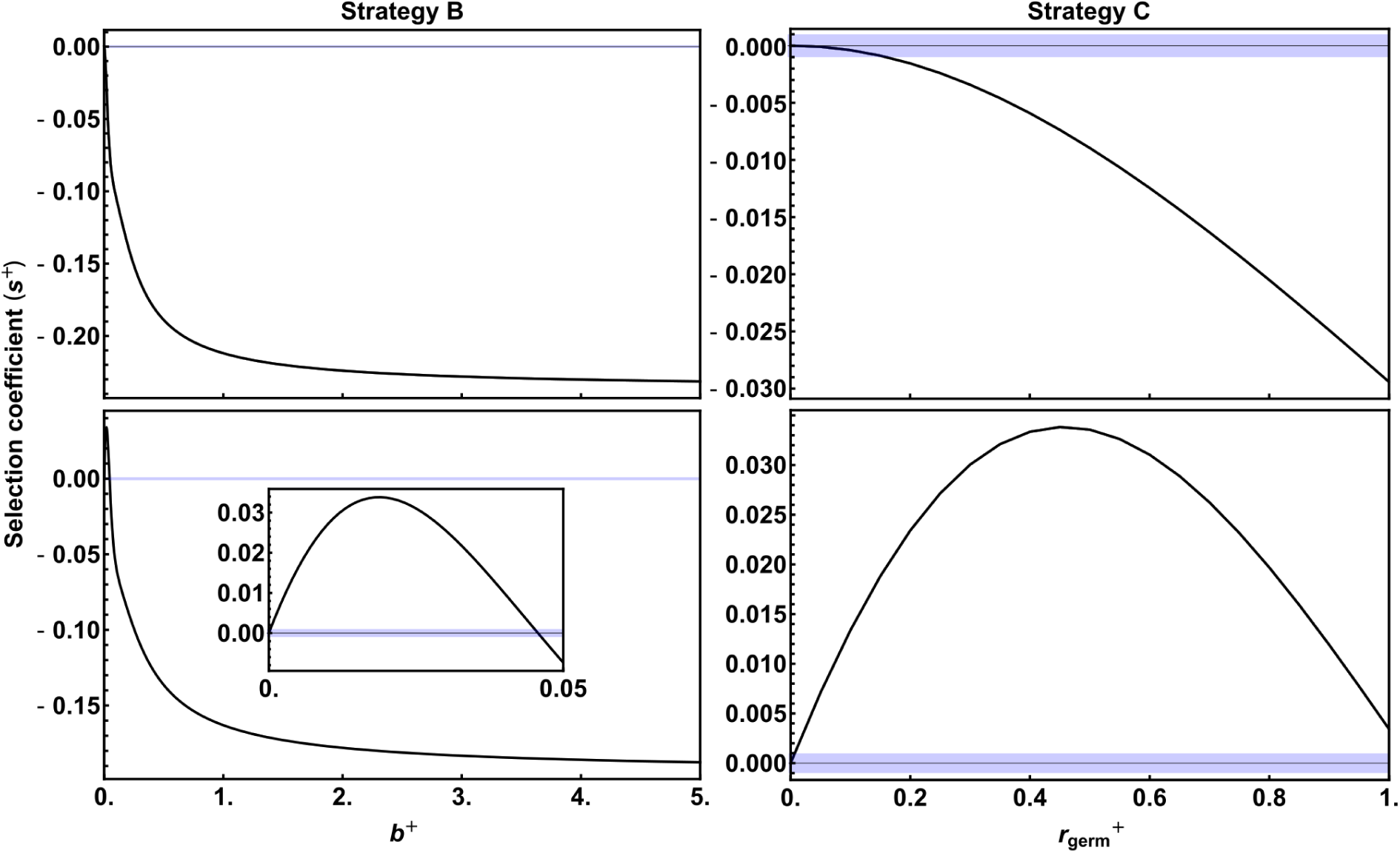
Selection for a fixed amplification rate (*b*^+^ > 0; strategy B in left panels) and transmission of sRNA transcripts (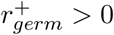; strategy C in right panels) in a resident population of strategy A (*b*^−^ = 0 and 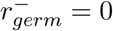). Upper panels show default dynamics (transcription rate optimal for strategy A), and bottom panels show dynamics slowed by a factor 0.75. Strategies A-C do not have a plastic response to the environment and so the curves are insensitive to the order of environments encountered (*p*_*ϵ*_). Blue shaded regions denote weak selection coefficients (| *s*^+^ |≤ 0.001).

Because we assume that strategy A has an sRNA transcription rate that is the best possible fixed strategy for the environmental conditions experienced, strategies B and C can only increase sRNA levels beyond the already optimal level produced via transcription and therefore cannot be fitter than strategy A. However, if the system is constrained such that sRNA abundance is not able to reach the optimal sRNA level within an individual generation, sRNA amplification (*b*^+^ > 0) and/or transmission 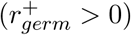 can become selectively advantageous.

To explore such constraints, we investigate a case where the dynamics are slowed by a factor 0.75 (multiplying *b, d, m*, and *µ* by 0.75), in which case the same steady state level of sRNA would be reached but over a longer period of time (Figure S2). With slower dynamics, sRNA levels do not reach the optimal fixed level within one generation. In this case, we find an advantage to amplification of sRNA or transgenerational inheritance of sRNA (Figure 4; bottom panels).

Although not explored, we expect that sRNA amplification and inheritance would also be favored if fitness differences between the strategies early in life mattered, as well as at the adult stage where selection acts in our model. While increasing amplification allows sRNA levels to accumulate faster (Figure S2), only the transmission of sRNA transcripts from parents to offspring (*r*_*germ*_ > 0) allows high levels of sRNA to be reached early in life (Figure 3), buffering juveniles from stressful environments.

### Evolution of environmentally responsive sRNA amplification and inheritance

We next examine whether a mutant capable of responding to current environmental conditions by changing the amplification rate *b* (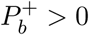; strategies D-F) could invade a resident population of non-amplifying, non-plastic individuals (strategy A; *b* = 0 and 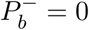). Recall that we focus on responsive strategies that increase the amplification rate in stressful environments (adaptive phenotypic plasticity; 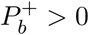 in see Equations 3-6).

We first consider mutants only capable of somatic changes in amplification rate based on the current environment (strategy D). Because such mutants increase the production of sRNA specifically when the environment is stressful, strategy D mutants can be selectively advantageous (*s*^+^ > 0) compared to the resident strategy A (Figure 5; red curves). Because the response is restricted to the soma, the similarity in environments experienced by parents and offspring does not affect selection for the plastic strategy D (red curves are insensitive to changes in *p*_*ϵ*_ across the panels in Figure 5). The selective benefit of strategy D is strongest at intermediate degrees of responsiveness to the environment 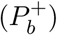, reflecting a balance between the benefits of having some plasticity and the costs of responding precisely.

**Figure 5:**
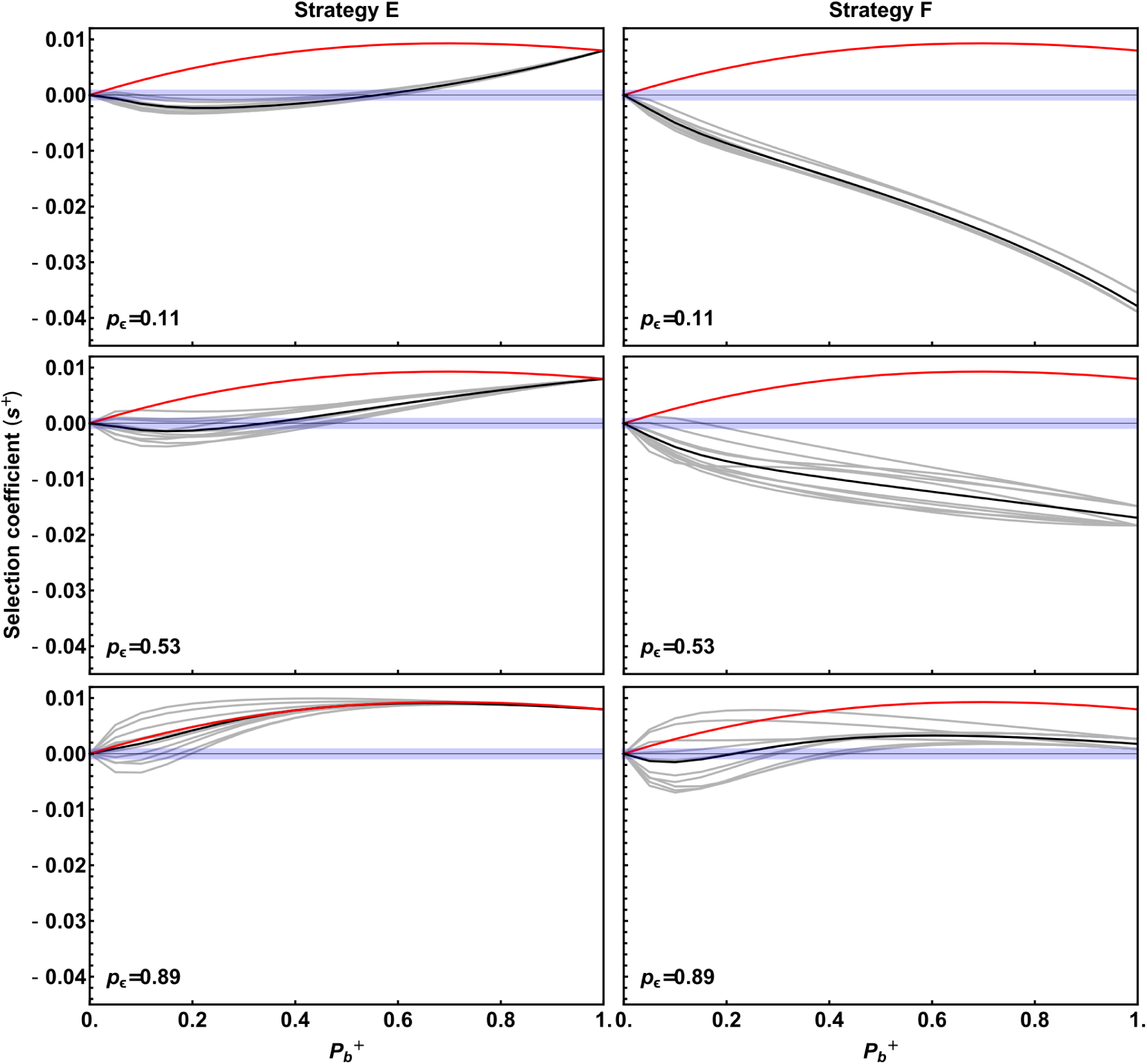
Selection on strategies that amplify sRNA in response to environmental conditions (strategies D-F). Red curves represent strategy D. Black curves represent the mean selection coefficient across different realizations (gray curves) of environmental scenarios with a given probability that parents and offspring encounter the same environment (*p*_*ϵ*_). Blue shaded regions denote weak selection coefficients (| *s*^+^ |≤ 0.001).

Selection for plastic sRNA amplification depends strongly on the cost of plasticity. Increasing this cost from *C*_*b*_ = 2*C*_*n*_ to *C*_*b*_ = 10*C*_*n*_ eliminated the benefits of strategy D under the default parameters (Figure S4). Essentially, because we allow strategy A to evolve first towards its optimum in the absence of plasticity, plastic sRNA amplification is favored only if it involves sufficiently low costs.

When plastic changes in the amplification rate can be transmitted to offspring, the selective advantage of mutants depends on the similarity of environments encountered by parents and offspring (*p*_*ϵ*_) and whether plastic sRNA amplification affects both the soma and germline (strategy E) or only the germline (strategy F) as illustrated in Figure 5. With transgenerational inheritance (strategies E and F), the exact order in which environments are encountered now influences the strength of selection (gray lines in Figure 5), even holding constant both the array of environments (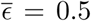 with *ϵ* ∈ {0.1, 0.9}) and the level of environmental similarity (*p*_*ϵ*_). Again, only when costs of plasticity are sufficiently low (small *C*_*b*_ *· P*_*b*_) does selection favor plastic strategies E and F over the wildtype (compare Figures 5 and S4).

Having both a plastic somatic and germline response (strategy E) is particularly favored when the environment remains the same for several generations (high *p*_*ϵ*_). By contrast, a plastic response restricted to the germline (strategy F) is never as strongly advantageous, because the restriction of amplification changes to the germline provides a fitness benefit only when the current and parental environments are both stressful. As a consequence, strategy F is rarely favored over the fixed wildtype strategy A when the environment varies frequently between parents and offspring (*p*_*ϵ*_ = 0.11, 0.53).

Even when fitter than strategy A (Figure 5), plastic mutants allowing the amplification rate to be transmitted across generations (strategies E and F, gray and black curves) are generally less fit than strategy D that has only a somatic response to the current environment (red curves). Again, we hypothesize that this is due to our choice of default parameters, which allow sRNA levels to reach optimal levels within a generation for strategy A, limiting the benefits of the “headstart” that could be provided to offspring in stressful environments that inherit a higher amplification rate from their parents. Again, if we slow down the dynamics (multiplying the within-generation parameters *b, d, m*, and *µ* by 0.75), transgenerational inheritance is much more commonly favored (Figure 6), especially if plasticity and its associated costs are low (small *C*_*b*_ *· P*_*b*_).

**Figure 6:**
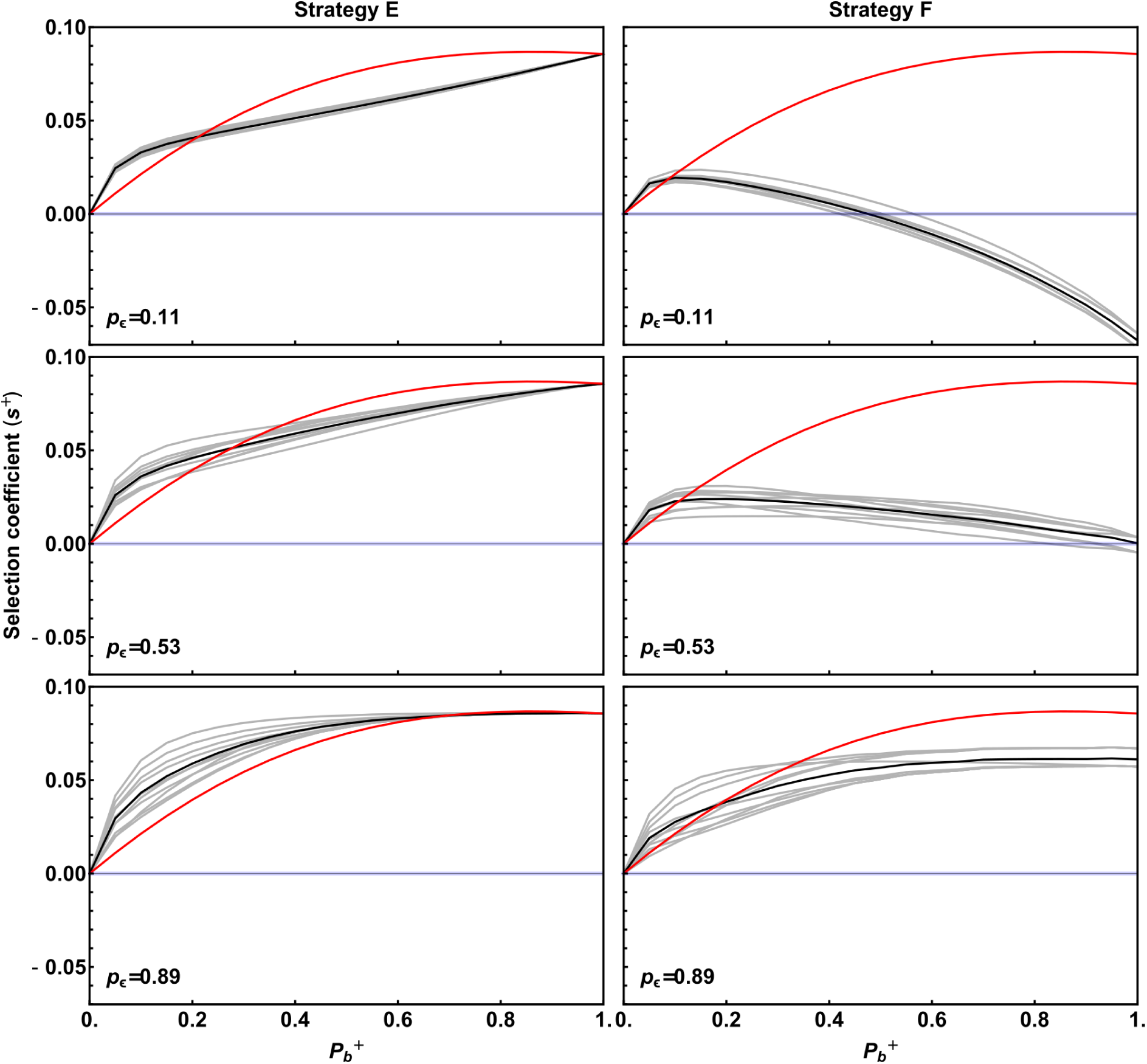
Selection on strategies that amplify sRNA in response to environmental conditions (strategies D, E and F), when the dynamics of sRNA within an individual are slowed to 0.75 times the original speed, preventing sRNA levels from reaching optimal levels within one generation. See Figure 5 for additional details.

Finally, we explore the selective advantage of mutations altering the germline transmission of sRNA in the context of plastic sRNA amplification by determining the full two-dimensional fitness landscape with respect to both plasticity in amplification rate 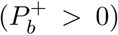 and sRNA transmission 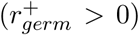. For ease of presentation, we randomly selected one environmental scenario with low environmental similarity (*p*_*E*_ = 0.11) and one scenario with high environmental similarity (*p*_*ϵ*_ = 0.89) to illustrate the fitness surface (Figure S5). For these parameters, the highest fitness strategies never involve inheritance of sRNA transcripts (the fitness maxima illustrated by black points in Figure S5 occur at 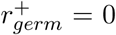). Thus, as we found with strategy C, plasticity does not inherently favor the transmission of sRNA transcripts when the within-generation dynamics are fast enough for sRNA levels to reequilibrate by the time that selection acts. We predict that if the wildtype is constrained (e.g., with slow sRNA dynamics within a generation) or selection acts at earlier stages in the life cycle, then direct sRNA transmission would be favored to buffer juveniles from stressful environments when parental and offspring environments are similar (high *p*_*ϵ*_).

## Discussion

Modeling the dynamics of sRNA that buffer the organism against stressful environments, we find that mechanisms that allow sRNA levels to be amplified under stressful conditions are favored unless the costs of plasticity exceed the benefits (strategy D; Figure 5). By contrast, the transmission of sRNA amplification rates across generations via non-genetic inheritance of sRNA templates, amplification agents, or epigenetic up-regulation of those agents (strategies E or F) is selectively favored over somatic plasticity (strategy D) only if the optimal sRNA level cannot be reached by the adult stage at which selection acted in our model (Figure 6). Similarly, transmitting sRNA transcripts directly to offspring (strategy C) did not increase fitness (Figure 4 and S5) over the wildtype (strategy A) unless the optimal sRNA level could not be reached before selection acted. Transmission of either sRNA amplification rates or sRNA transcripts from parents to offspring is therefore most likely to evolve when constraints prevent sRNA levels from rising to high levels within a single generation and when plastic responses are slow and unable to provide immediate benefits (strategies D and E unavailable).

Based on the dynamics of sRNA levels within a single generation (Figure 3), the transmission of sRNA transcripts is particularly likely to be favored when selection acts at juvenile stages, when transcript levels have yet to reach high levels in the absence of transgenerational inheritance, although we did not directly explore this conjecture as fitness depended only on adult levels of sRNA in our model (Equation 7).

Previous theoretical studies have found similar results regarding the benefits of non-genetic inheritance (e.g., maternal effects) under various environmental conditions. Kuijper *et al*. [71] showed that phenotypic memory in the form of a positive correlation between parental and offspring phenotype is favored when environmental conditions are relatively stable. As the rate of change of environmental conditions increase, that correlation decreases and selection favors less faithful transmission of phenotypes. When maternal effects are considered from a multivariate perspective, selection favors a positive correlation between the multivariate phenotypes of the offspring and the mother when the rate of environmental fluctuations is low, but a negative correlation in rapidly changing environments [72]. Furthermore, informative maternal effects were shown to be beneficial when juvenile cues are inaccurate, transmission of maternal cues are accurate, and the environment is highly stable [73]. Although the transmission of phenotypes from parent to offspring may be beneficial under certain environmental conditions (mentioned above), parents and offspring may have conflicting interests that can affect the evolution of information transfer from parents to offspring. Kuijper *et al*. [74] explored the effect of such parent-offspring conflicts and showed that, in many cases, it causes a partial or complete breakdown of informative maternal effects, which may explain the apparent weakness of transgenerational plasticity in nature.

When fitness is affected by environmental conditions, different systems of phenotypic determination can evolve depending on the accuracy of genetic versus environmental cues. When genetic cues are accurate and environmental cues are inaccurate, phenotypic determination based on genetic polymorphism is likely to evolve. However, when genetic cues are inaccurate and environmental cues are accurate, phenotypic plasticity is more likely to evolve [75; 76; 77]. These outcomes have important consequences for the interpretation of our results, in particular for strategies E and F, where transgenerational plasticity is present. In highly autocorrelated environments (high *p*_*ϵ*_), strategies E and F parents have an accurate environmental cue and can benefit their offspring by transmitting their sRNA amplification rate to gametes, which is similar to results from previous models on maternal effects without the amplification dynamics considered here [78]. With weakly or negatively autocorrelated environments, however, the sRNA amplification rate transmitted by the parents is frequently detrimental to the offspring, thereby favoring somatic plasticity without transmission (strategy D over strategies E and F), unless transgenerational inheritance is needed to reach high sRNA levels (compare Figures 5 and 6 for fast and slow dynamics, respectively). Additionally, previous results have concluded that maternal effects are more strongly favored when plasticity is limited [78], which is consistent with our results that non-genetic inheritance of sRNA amplification rates (strategy F) is typically not favored if a highly plastic strategy is available (strategy D).

Unlike previous theoretical studies, however, our study provides solid, clear predictions for the well-known molecular mechanism of sRNA-mediated phenotypic plasticity, sRNA amplification, and non-genetic inheritance. Additionally, we provide a set of possible strategies (strategies B-F) that diverge from an ancestral state (strategy A) in specific molecular mechanisms (Figure and predict the evolutionary fate of such mutations under a series of different environmental scenarios. Our model thus provides a valuable reference for future empirical studies testing the effect of sRNA-related mutations and the effect of environmental stress on such mutants.

The predicted evolutionary response of sRNA production and transmission to changing environments obtained from our model is in line with recent empirical observations in the nematode *C. remanei* [in press, 79]. Nematodes were exposed to four different environments including stable at a control (20°C) or high temperature (25°C), or in slowly or rapidly changing environments. Lines exposed to stable or slowly changing environments showed a strong positive maternal effect on reproductive output resulting in increased offspring production, whereas lines maintained in a rapidly changing environment showed a reduced maternal effect on off-spring production. This finding is consistent with our results and others [73; 80; 71] that the transgenerational transmission of non-genetic information (parental effect) about environmental conditions is more beneficial when environmental conditions are similar between parents and offspring. The mechanism underlying these observations is currently unknown, however, and identifying these will be an interesting next step.

sRNAs are known to play an important role in mediating phenotypic plasticity across different taxonomic groups and are believed to be part of an adaptive response to fluctuating environmental conditions. In *C. elegans*, for example, developmental arrest caused by starvation leads to a regulation of endogenous siRNA and their mRNA targets primarily associated with nutrient storage [49]. In the thale cress, *Arabidopsis thaliana*, siRNA production is induced by salt stress and reduces the expression of a proline catabolic gene, leading to proline accumulation, which is important for salt tolerance [81; 82]. Further evidence comes mostly from studies of responses to biotic stress, namely pathogenic infections in both plants and animals [83]. In plants, the role of sRNAs in silencing pathogen gene expression has been established and seems to be widespread [84; 85]. In *A. thaliana*, cells infected with the fungal pathogen *Botrytis cinerea* secrete vesicles that deliver sRNAs into the fungus, inducing silencing of the fungal genes associated with pathogenicity [86]. In some cases, pathogen counter-defense can evolve leading to a molecular arms race between the hosts’ silencing mechanisms and the pathogens’ evasion strategies [87]. Similarly, *C. elegans* uses sRNA interference as an immune response to artificial viral infections, with RNAi-defective worms displaying aggravated infections while RNAi-enhanced worms inhibit the production of infectious progeny virus [88; 89]. This reduction of pathogenic infections caused by the production of sRNAs supports the idea that sRNAs play a role in adaptation to novel environments. Furthermore, pathogenic infections are typically localized in time and space, providing the type of autocorrelated environments that we find favor transgenerational inheritance of sRNA amplification rates.

sRNAs are also known to have a number of functions in the germline and can be transmitted between generations. In *Drosophila*, piRNAs have been shown to protect the germline against the genomic replication of transposable elements by targeting transposon transcripts [90; 91]. In addition, inherited germline-derived piRNAs affect the deposition of histone marks and initiation of primary piRNA biogenesis in the offspring and thereby act as an epigenetic memory across individual generations [92; 93]. Similarly, in *C. elegans*, worms infected with different viruses (e.g., flock house virus, Orsay virus) not only develop an siRNA-based antiviral response to silence viral RNAs and stop the infection, but they also pass this response on to their offspring [48; 94]. This transgenerational immunogenic memory works as a form of inherited vaccination against future infections [95]. Additionally, the inheritance of sRNAs (piRNAs and siRNAs in particular) allows the transmission of information about previous exogenous viral RNA infections in *C. elegans* and information about endogenous stress-related processes, like starvation. Worms derived from starved great-grandparents and kept fed for three individual generations showed a siRNA response to their previous ancestors’ starvation experience, with differentially expressed siRNAs targeting genes involved in nutrition. The transmission of sRNA levels was shown to persist for up to four generations without further stimulation [49].

While the role of sRNAs in protecting the soma and germline from environmental stressors and the transmission of sRNAs across generations are undeniable, the adaptive value of the transmission of sRNAs across generations is debated. However, recent findings in *C. remanei* suggest that non-genetic inheritance may be an adaptive response to unstable environments [in press, 79]. The evolution of plastic amplification rates for sRNAs might provide hosts with an edge in the race against everchanging viruses and other disease agents. Indeed, disease agents may sometimes supply the sRNA whose amplification has been co-opted for host defense [96]. Similarly, abiotic environmental factors might drive the evolution of plastic amplification processes to help organisms maintain their physiological state (homeostasis) when exposed to adverse environmental conditions. Here, we provide a proof-of-concept model for the role of environmental fluctuations in the adaptive evolution of sRNAs, considering both plastic amplification in response to current conditions and the potential for inheritance of sRNA transcripts or amplification rates. Future empirical studies are needed to explore the fitness advantages of transgenerational transmission of sRNAs as well as the mechanisms by which plastic sRNA amplification is transmitted from parents to offspring through epigenetic marks or cytoplasmic amplification agents.

## Acknowledgements

WTAFS was funded by a grant from the Sven and Lilly Lawski Foundation. SPO was funded by a Natural Sciences and Engineering Research Council of Canada grant (RGPIN-2016-03711). SI was funded by grants from the European Research Council (ERC Starting Grant HapSelA-336633), the Human Frontier Science Program (HFSP R0025/2015) and the Knut and Alice Wallenberg Foundation.

## Competing interests

The authors declare that they have no conflict of interest.

## Author contributions

WTAFS further developed the numerical model and ran the simulations. SPO and SI conceived the initial numerical model. All authors contributed to the design and implementation of the research, the discussion of the results and the writing of the manuscript.

## Supplementary Information

**Figure S1:**
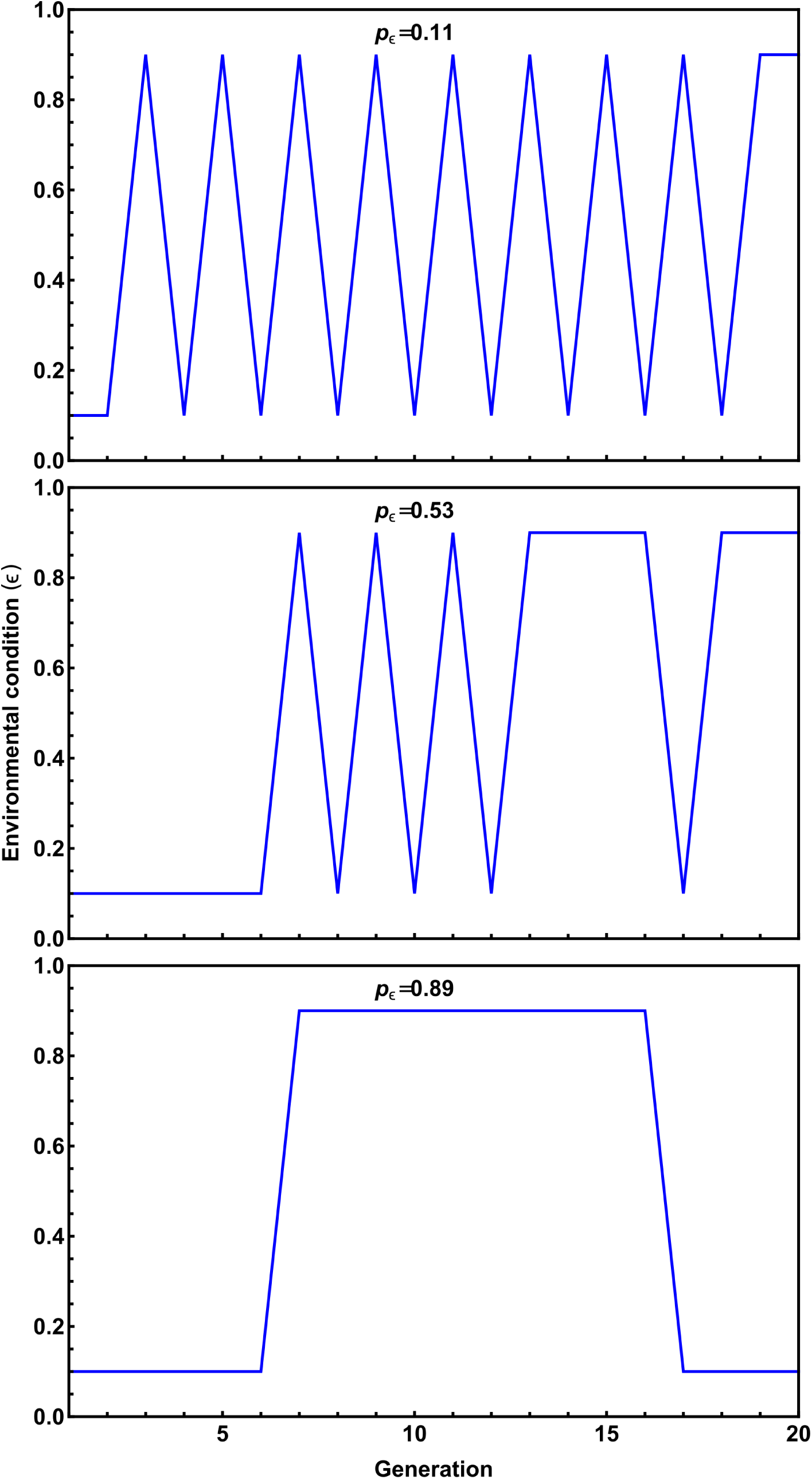
Examples of environmental scenarios with different levels of similarity between the environment of parents and offspring (*p*_*ϵ*_). Note that we held the number of relaxed (*ϵ* = 0.1) and stressful (*ϵ* = 0.9) environments constant (50% each) to allow us to optimize the system in the absence of epigenetic inheritance.

**Figure S2:**
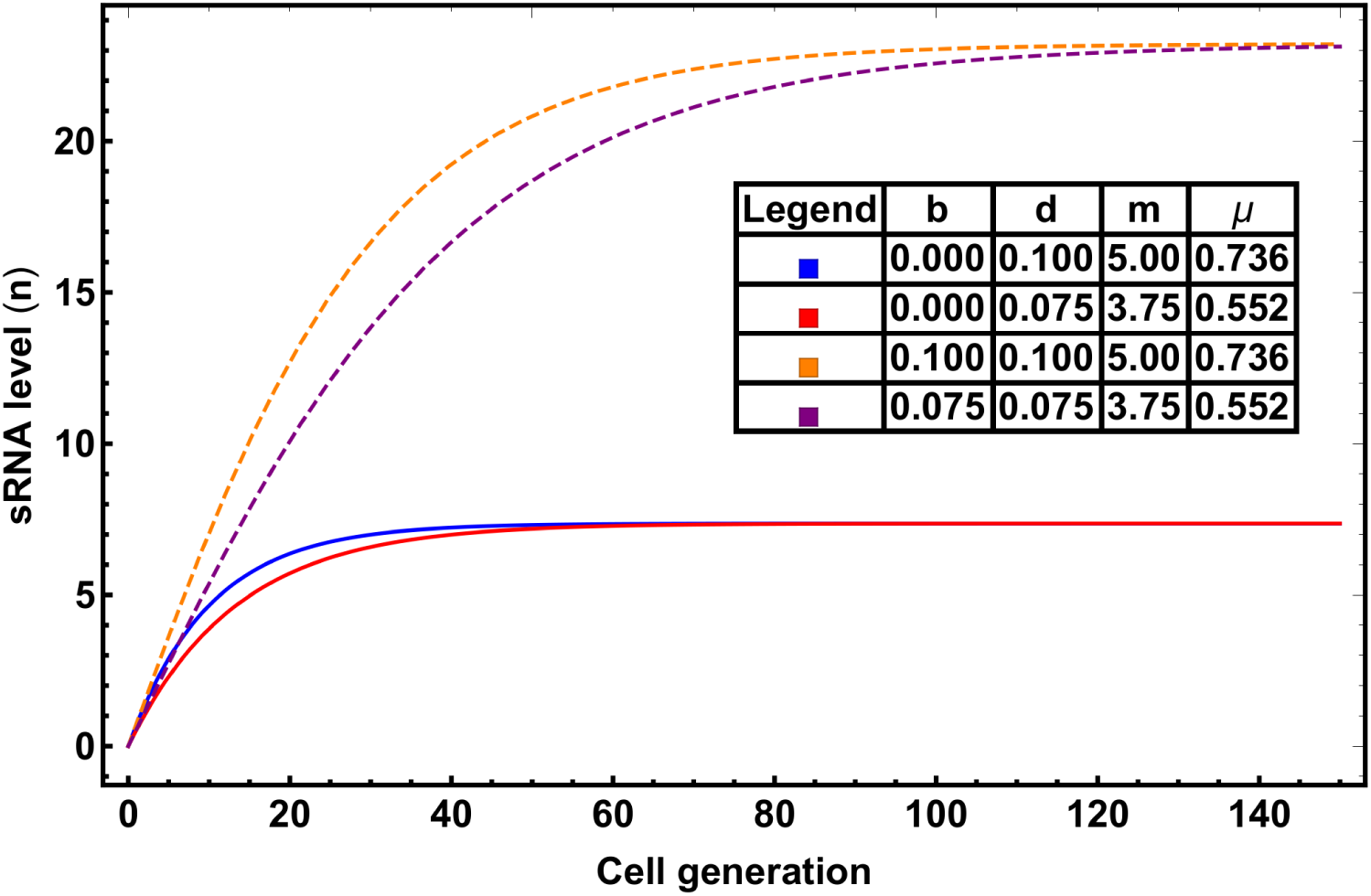
Dynamics of individual sRNA production across cell divisions under different biological conditions (parameter values). Amounts of sRNAs (*n*) reach their steady state values over time, measured in cell divisions. Solid lines show dynamics in a wild type individual (with *b* = 0; fast and slow dynamics), and dashed lines show dynamics in an amplifying mutant (*b* > 0; fast and slow dynamics).

**Figure S3:**
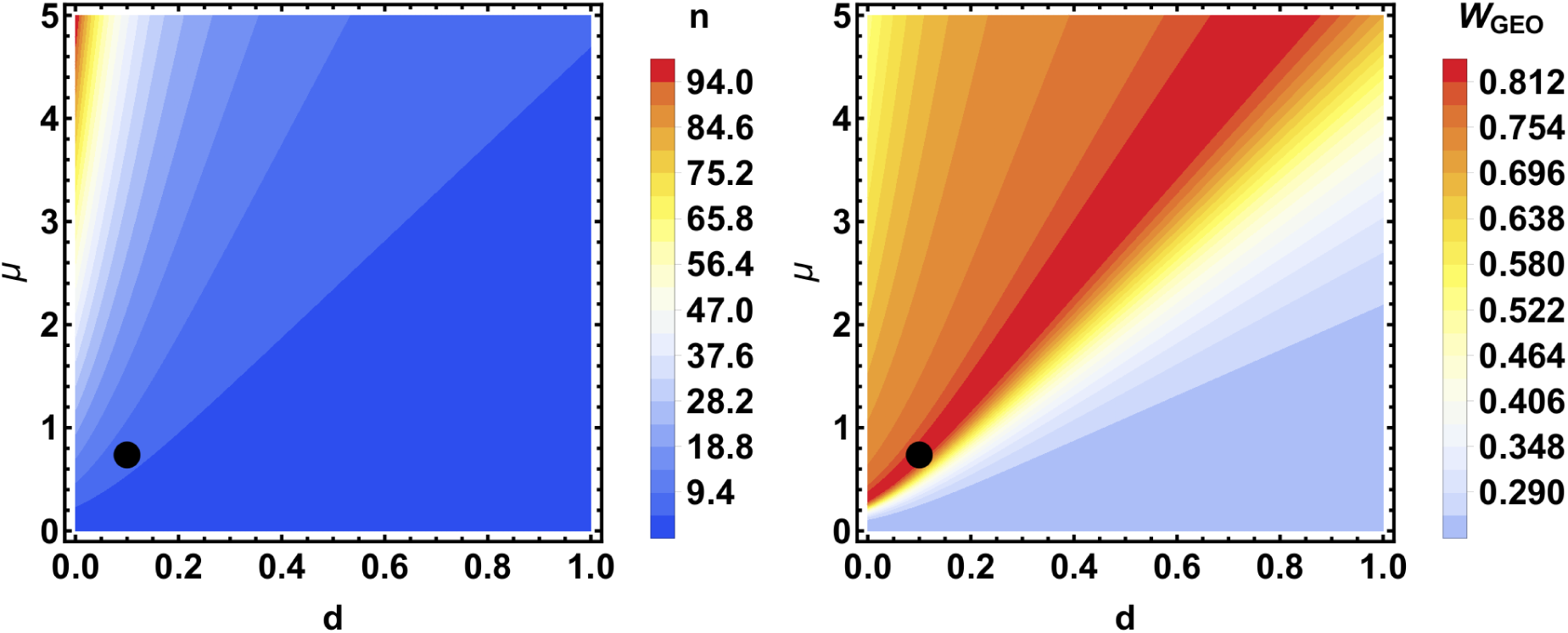
Production of sRNAs (*n*) given different transcription and degradation rates, under wildtype conditions (*b*^−^ = 0, 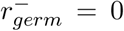 and 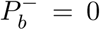; left), and its geometric mean fitness (*W*_*GEO*_) under fluctuating environmental conditions (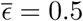; right). The black circle indicates the intrinsic degradation rate (*d* = 0.1) and transcription rate (*µ*^−^ = 0.736) that was optimal with 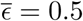 and was used across the study.

**Figure S4:**
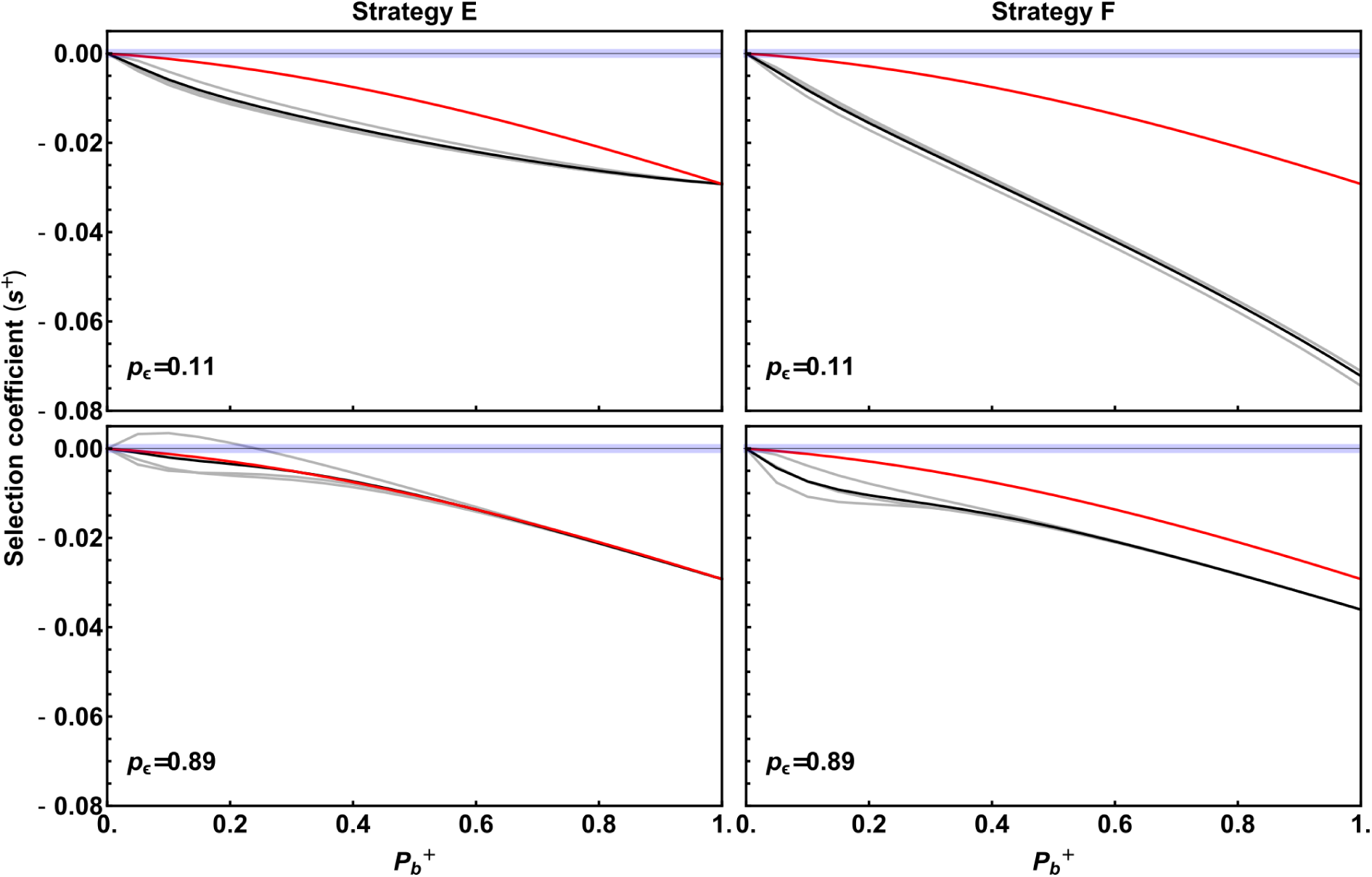
Invasiveness of strategies D, E and F when the cost of plasticity (*C*_*p*_) is ten times higher than the cost of sRNA production (*C*_*n*_). See Figure 5 for additional details.

**Figure S5:**
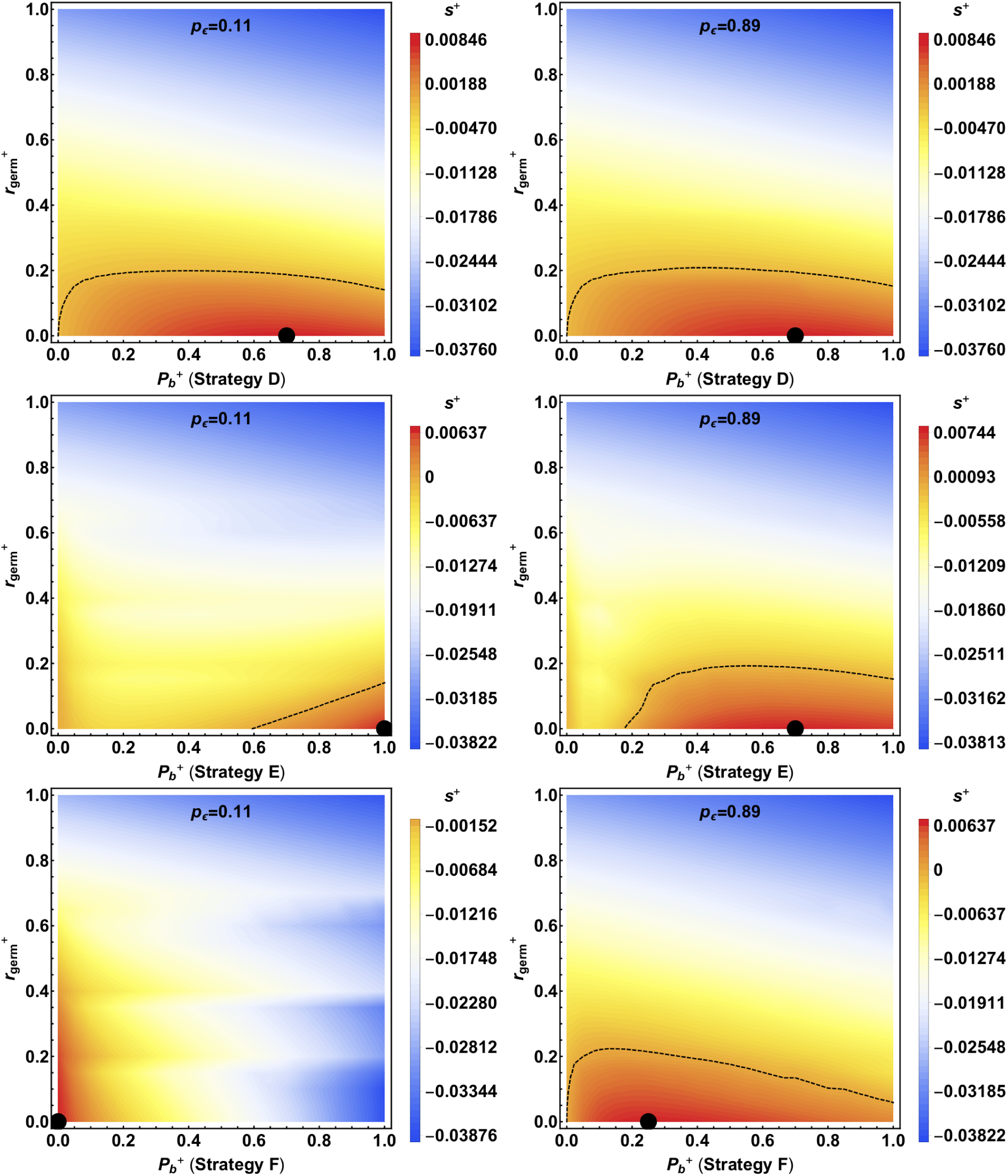
Invasiveness of strategies D, E and F (*P*_*b*_ > 0) coupled with transmission of sRNA transcrips to offspring (*r*_*germ*_ > 0). Black points indicate the highest geometric mean fitness relative to strategy A (selection coefficient). Dashed lines indicate *s*^+^ = 0, separating selectively beneficial parameter combinations from detrimental combinations.

